# Adaptive laboratory evolution of a yeast co-culture chassis for modular bioproduction

**DOI:** 10.64898/2026.09.06.747466

**Authors:** Diego Ruiz-Sanchis, Javiera López-Salinas, Cinzia Klemm, María Gallego García, Rodrigo Ledesma-Amaro

**Affiliations:** Department of Bioengineering, Imperial College London, London SW7 2AZ, UK; Bezos Centre for Sustainable Protein, Imperial College London, London SW7 2AZ, UK; UKRI Engineering Biology Mission Hub on Microbial Food, Imperial College London, London SW7 2AZ, UK; Advanced Biofuels and Bioproducts Unit, Department of Energy, CIEMAT, Avenue Complutense 40, 28040 Madrid, Spain; Alcalá de Henares University, Alcalá de Henares, Spain

## Abstract

Synthetic microbial consortia have the potential to bring novel architectures to biotechnological processes. They enable more flexible and efficient bioprocesses by reducing metabolic burden and supporting division of labour. Obligately mutualistic cross-feeding has been used to stabilise the population composition of artificially assembled consortia. However, many current strategies rely on the cross-feeding of metabolites with limited exchange rates, and this imposes a growth burden on the consortium members. Here, we used adaptive laboratory evolution (ALE) to address this growth bottleneck and create an optimised co-culture chassis to host bioproduction functions. Transcriptome analysis revealed how ALE alleviated stress responses associated with nutritional restrictions in the non-evolved cross-feeding system. Using a case-study split bioproduction pathway, we demonstrated that improvements in the chassis growth performance resulted in production improvements, surpassing the performance of a monoculture implementation by 1.5-fold. Our results show the potential of ALE to optimise the cross-feeding layer of yeast co-cultures, and how this enables the efficient implementation of a bioproduction process.

## Introduction

Synthetic microbial consortia have emerged as a promising tool in the field of biotechnology, as they enable the design of modular processes to produce food ingredients^1^, pharmaceuticals^2^, biomaterials^3,4^ or fuels^5^. Synthetic consortia translate the principles and inherent properties of natural microbial communities to technological challenges: distributing bioprocesses across microbial communities can increase productivity by dividing cellular and metabolic burden^6,7^, preventing metabolic promiscuity by compartmentalising biochemical reactions^8,9^, as well as simplifying pathway optimisation by allowing modular interventions^10^. However, the outcome of these applications is highly dependent on the relative population size of each strain in the community, which leads to major differences in productivity depending on the inoculation ratio and the relative growth rate of community members^11,12^.

Cross-feeding interactions are widespread in natural microbial communities^13^ and have been used as a mechanism to increase the population robustness and stability of synthetic consortia^14,15^. This is particularly relevant for metabolic engineering implementations, where the relative fitness of strains may significantly differ due to differences in burden caused by heterologous gene expression^16^. However, engineering cross-feeding remains challenging in yeast cell factories, because the mechanisms governing metabolite secretion are still not fully understood^17,18^. In previous works, synthetic cross-feeding has been achieved by introducing feedback-resistant alleles to support the overproduction and overflow of amino acids and nucleotide bases^19,20^. More recently, specific pairs of *Saccharomyces cerevisiae* auxotrophs for this type of metabolites were observed to engage in synergistic growth without further genetic modification, challenging prior assumptions that this was not feasible^21^. Despite these advances, current yeast co-cultures that rely on the exchange of amino acids and nucleotide bases for population control, typically exhibit slow growth and their viability is limited to specific inoculation regimes^20–22^.

Adaptive laboratory evolution (ALE) is well-suited to address engineering challenges whose underlying mechanisms are poorly understood. In metabolic engineering, ALE has been applied to enhance titres and productivity^23^ or to improve growth on non-native substrates^24,25^. ALE has also been applied to microbial consortia: co-evolution experiments have provided insights into community stability and the emergence of ecological interactions^26,27^, improved overall community growth performance^28^ and enhanced metabolite secretion^29^. However, in the absence of a dedicated selection strategy, ALE of microbial consortia can result in the loss of the heterologous bioproduction function that is hosted, as its associated production burden creates a negative selection pressure^30^.

In this study, we sought to examine how ALE can be used to improve the cooperative ecology of a *S. cerevisiae* co-culture with the goal of improving its capacity to ultimately host a biomanufacturing process. We subjected three different obligately mutualistic cross-feeding co-cultures to ALE, and isolated evolved clones where the initial growth limitation had been significantly alleviated. We used transcriptomics to contextualise these adaptations. Finally, we introduced a heterologous bioproduction function in this chassis and studied how it was affected by the changes that ALE had elicited, finding that the evolved chassis led to higher production levels and exceeding the performance of a monoculture.

## Results

### The outcome of experimental evolution varies between co-cultures depending on the nature of cross-feeding

We wanted to explore the potential of ALE to improve the growth of obligately mutualistic cross-feeding co-cultures. To do this, we chose three co-cultures that had been previously characterised in our lab. These are composed of two auxotrophic strains that rely on the reciprocal exchange of amino acids or nucleotide bases for survival. One of the co-cultures was identified in the high-throughput screening by Aulakh *et al*. ^21^ (*his2*Δ:*met3*Δ, from now on *his:met*), and two were chosen from the toolkit by Peng *et al*. ^20^ (*lys2*Δ-*ADE4*^op^: *ade8*Δ-*LYS21*^op^, from now on *ade:lys*; *lys2*Δ-*TRP2*^op^: *trp1*Δ-*LYS21*^op^, from now on *trp:lys*) (**Fig. 1a**).

**Figure 1.**
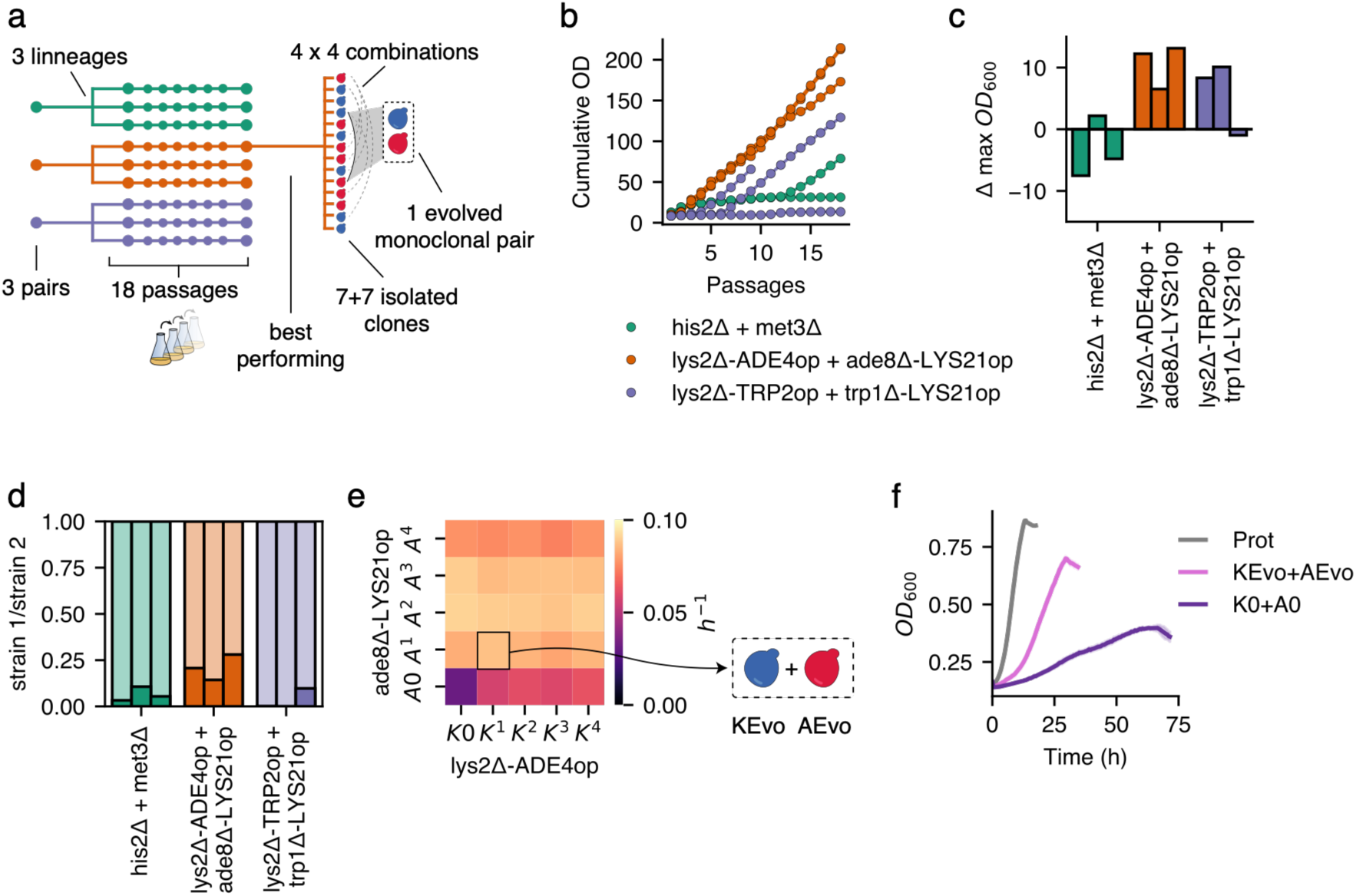
Adaptive laboratory evolution (ALE) of obligately mutualistic cross-feeding co-cultures. **a** Three cross-feeding cocultures were subject to serial passaging. Three lineages of each co-culture were subject to ALE in parallel, clones were isolated form the best performing one, combinatorially tested, and a single, monoclonal pair was chosen for further study. **b** Cumulative optical density (OD_600_) of each evolving lineage along 18 passages. **c** Difference in the saturation OD_600_ of each lineage between the final the second passage of the ALE course. **d** Population composition in the final passage of each lineage. Each bar represents a single lineage. The bottom, coloured segment of each bar represents the fraction of the total population corresponding to the first strain listed in each x-axis label (“strain 1 + strain 2”). **e** Maximum growth rate of each combination of clones isolated from the lys2Δ-ADE4op + ade8Δ-LYS21op co-culture. K0 is the ancestral lys2Δ-ADE4op strain, K^1-4^ are evolved clones of this same strain; A0 is the ancestral ade8Δ-LYS21op strain, A^1-4^ are its evolved clones. Clones A^1^ and K^1^ will be referred to as AEvo and KEvo from now on. Each tile represents the average value of two replicates. **f** Growth of the ancestral lys2Δ-ADE4op + ade8Δ-LYS21op co-culture (K0+A0), a co-culture between the clones KEvo and AEvo, compared to a prototrophic monoculture (Prot). Solid lines represent the average of three replicates, and the shaded area around it depicts standard deviation.

Three replicates of each co-culture were subjected to serial passaging in minimal medium for 18 rounds, and their growth and population composition were tracked over time. Sustained growth across all 18 passages was observed in all three *ade:lys* lineages, but only in one of the *his:met* and one of the *lys:trp* lineages (**Fig. 1b**). To study the effect of ALE on carrying capacity, we compared the maximum optical density (OD) reached by the co-cultures at the second passage and at the last passage of the ALE course. All three *ade:lys* lineages showed an improvement in saturation OD of more than 6 units, with the best-performing lineage reaching a 4.6-fold relative increase. Conversely, only two of the three *lys:trp* lineages improved their carrying capacity, while very little improvement or even a decrease in this metric was measured in the *his:met* co-culture (**Fig. 1c**). Population composition analysis of the co-cultures in their final passage revealed a consistent persistence of both strains in the *ade:lys* co-culture across all three lineages, averaging around a 20:80 ratio. Coexistence was also maintained in all *his:met* co-cultures, but at more off-centre ratios. Interestingly, the two lys:trp lineages that showed an improvement in carrying capacity were completely taken over by the lysine-overproducing strain (**Fig. 1d**).

Based on the metrics above, we chose the *ade:lys* co-culture to further develop a bioproduction platform. We isolated random clones from this population and benchmarked their growth. Four clones of strain *lys2*Δ-ADE4^op^ (K^1^, K^2^, K^3^, K^4^) were combinatorially tested with four *ade8*Δ-LYS21^op^ clones (A^1^, A^2^, A^3^, A^4^). We found that their growth rate (**Fig. 1e**) and carrying capacity (**Supplementary figure 1**) were homogeneous, except for combinations involving clone A^4^, which resulted in lower growth rate in all combinations (**Supplementary figure 2**). Interestingly, no significant differences were observed between co-cultures involving the non-evolved strain K0 and those containing the evolved clones K^1-4^, so long as the partnered strain was one of the evolved A^1-4^ clones (p=0.82 in Kruskal-Wallis test, **Supplementary figure 3**). This suggests that the fitness improvement of the evolved co-culture is primarily associated with adaptations in the A lineage. We selected the pair consisting of K^1^ and A^1^ for further study, which from now on will be referred to as KEvo and AEvo, respectively. As a result of this experimental evolution process, we isolated a co-culture with a significant improvement in growth parameters, more closely resembling the growth dynamics of a prototrophic monoculture (**Fig. 1f**).

### Cellular response to obligately mutualistic co-culturing before and after ALE

To further clarify the effects of cross-feeding and ALE on co-culture physiology and how this impacts bioproduction, we designed a strain-resolved transcriptomics experiment studying co-cultures of A0 and K0, AEvo and KEvo, as well as a prototrophic monoculture (Prot). Co-cultures were grown in split growth chambers divided by an exclusion membrane and strains were processed separately for RNA-Seq analysis (**Fig. 2 a**).

**Figure 2.**
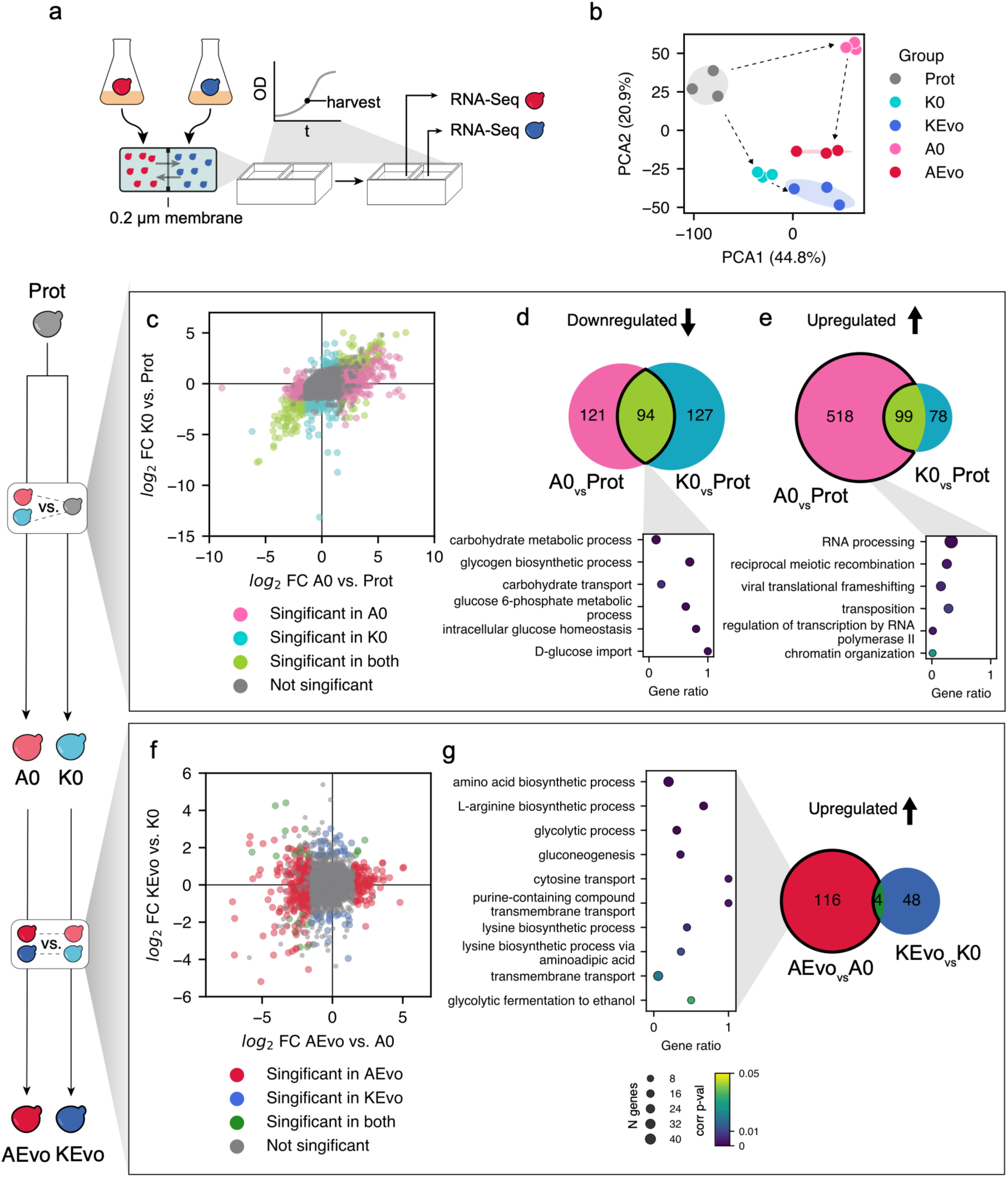
Transcriptome changes in the K0:A0 and KEvo:AEvo co-cultures. **a** Workflow for strain-resolved transcriptomics in a co-culture. Monocultures were washed and inoculated in either side of a split growth chamber separated by a 0.2 μm cell exclusion filter. Cells were harvested in mid-exponential phase and processed for RNA-Seq. **b** Principal component analysis of gene count data showing how A0 and K0 significantly diverge in their transcriptome, but this distance is minimised after ALE in AEvo and KEvo. **c, d, e** Differentially expressed genes as a result of entering an obligately mutualistic metabolism: comparison between A0 or K0 and a prototrophic monoculture (Prot). **c** Combined differential expression analysis showing log2 fold change in either strain along the axes. Colour palette shows significance threshold as per legend, defined as log2 fold change > 1.5 and padj < 0.05. **d** Intersection of downregulated genes in A0 and K0. 94 shared genes are highlighted and their GO enrichment results presented below. **e** Intersection of upregulated genes in A0 and K0. 518 genes are exclusive to A0, and their GO enrichment results presented below. Enrichment plots in d and e show ReviGo grouped terms. The dot diameter represents gene count, and the colour scale depicts the p-val as per the colour scale in panel g. **f, g** Differentially expressed genes resulting from the ALE process: comparison between AEvo and A0, and KEvo and K0. **f** Combined differential expression plot idem as in panel c. **h** Intersection of upregulated genes in KEvo and AEvo against their ancestral counterparts K0 and A0. GO enrichment of 116 genes upregulated exclusively in AEvo is shown below. Legends for corrected p-value colour scale and gene count in panels d, e and g are provided below panel g.

Our analysis focused on comparing the transcriptional response in two phases: first, when cells enter an obligately mutualistic metabolism by comparing a K0+A0 co-culture with a prototrophic monoculture (Prot) and, second, how ALE modified this response by comparing KEvo with K0 and AEvo with A0. Principal component analysis (PCA) of gene expression counts revealed two clearly distinct trajectories for either strain (**Fig. 2 b**). The initial transcriptional response of K0 and A0 entering co-culture was noticeably divergent, evidencing the impact of the nature of the cross-fed metabolite on the cellular response to co-culturing. Interestingly, ALE dampened the initial strain-specific response of A0 and reduced its disparity with K0: the distance between the KEvo and AEvo clusters was significantly reduced.

To identify which biological processes were part of the initial response, we performed differential gene expression analysis comparing K0 and A0 against a prototrophic monoculture (**Fig. 2 c**). A large set of differentially expressed genes (206 of 1006 DEGs, 20%) was shared across both groups, indicating that the initial response to co-culturing was only partially divergent. We further studied 94 DEGs which were downregulated in both A0 and K0 by over-representation analysis (ORA) for gene ontology (GO) enrichment of biological process terms. This revealed a shared downregulation of genes associated with carbohydrate import and metabolism, including hexo- and glucokinases, glucose transporters, and glycolytic genes (**Fig. 2 d**). This could suggest a slowdown of carbon utilisation and general metabolism as a response to nutritional stress from limited adenine and lysine cross-feeding, which is undesirable in a bioproduction context.

We also studied the subset of 518 genes that were upregulated in A0 but not in K0, seeking to explain the divergent component of the initial transcriptional shift of A0. Notably, we found a significant upregulation of genes involved in retrotransposition and associated processes (**Fig. 2 e**), which may reflect physiological responses associated with adenine limitation, as previously reported^32,33^.

We then turned to study the transcriptional changes that occurred as a result of adaptive evolution. This response was found to be very divergent, with only 25 of 536 unique DEGs (4.7%) shared between strains. Most transcriptional changes were found in the AEvo against A0 comparison, which accounted for 451 of 536 unique DEGs (84%) (**Fig. 2 f**). Among these, we found that the subset of genes upregulated exclusively in AEvo was enriched in carbohydrate metabolism enzymes, indicating a reversal of the initial downregulation response. Furthermore, we found an enrichment of amino acid biosynthesis genes, including the cross-fed metabolite lysine, but also arginine, and a significant enrichment of purine transporters, which responds to the need of K0 and KEvo strains to import adenine from the extracellular medium. Lastly, among the subset of downregulated genes in AEvo we found GO terms that we had previously associated with an activation of transposons, indicating that this response was alleviated by the action of ALE (**Supplementary figure 4**).

Whole-genome sequencing analysis provided further insights into some genetic changes underlying the transcriptome- and phenotype-level changes discussed above. Namely, we identified a large copy number variation of around 60 kb in chromosome V of AEvo (**Supplementary figure 5**). This segment contains 42 annotated ORFs, among which we found three genes coding for purine permeases: *FCY2*, *FCY21* and *FCY22*, which is consistent with the upregulation of purine transport found in the transcriptome analysis. This segment also contained the ORF *ARG56*, coding for the enzyme catalysing the third step in arginine biosynthesis, which was also found to be significantly upregulated in the transcriptomics dataset (**Supplementary figure 6**).

### Emergent phenotypes underlying improved growth in the evolved co-culture

Further study of the isolated KEvo+AEvo co-culture revealed several growth-related phenotypes that coevolved during the course of ALE. First, we noticed that lysine supplementation alone was sufficient to support the growth of a K0 monoculture in minimal media, but not KEvo (**Fig. 3 a**): a second amino acid had to be supplemented along with lysine to rescue its growth. We observed this effect with arginine supplementation and other 14 amino acids and nitrogen metabolites from a range of 22 we tested (**Supplementary figure 7**). Interestingly, this growth defect was not observed when KEvo was co-cultured with AEvo. One possible explanation is that ALE increased the availability of these nitrogen metabolites within the co-culture, which is consistent with the copy number expansion and transcriptional upregulation of *ARG56* observed in AEvo (**Supplementary figure 5**). In all further experiments that involved the growth of K0 or KEvo in a monoculture, arginine was supplemented to the medium.

**Figure 3.**
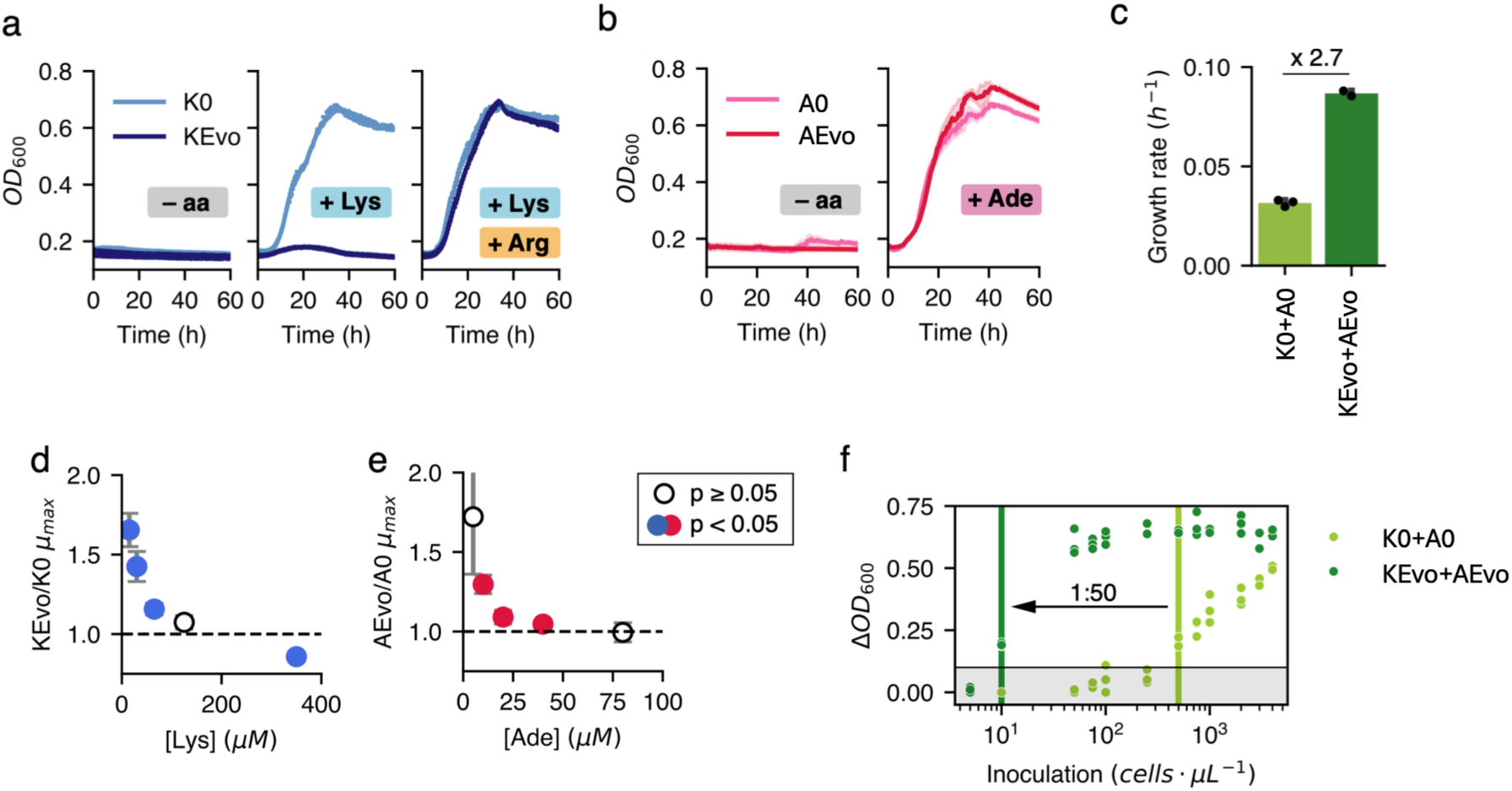
Growth-related phenotypes in the evolved coculture. **a** Growth of the ancestral lysine auxotroph (K0) and its evolved isolate (KEvo) in minimal medium (– aa), minimal medium supplemented either with lysine (+ Lys) or with lysine and arginine (+ Lys, + Arg). **b** Growth of the ancestral (A0) and evolved (AEvo) adenine auxotroph in minimal medium (– aa) and minimal medium supplemented with adenine (+ Ade). **c** Maximum growth rate of the ancestral (K0+A0) and evolved (KEvo+AEvo) cocultures. **d** Fold change in saturation optical density (OD_max_) in monocultures of the evolved blue auxotroph (KEvo) compared to its non-evolved ancestor (K0) in minimal medium supplemented with varying concentrations of lysine. Filled makers indicate p < 0.05 in Welch’s t-test. **e** Fold change in saturation optical density between the evolved and ancestral adenine auxotroph, at varying adenine concentrations. Filled makers indicate p < 0.05 in Welch’s t-test. **f** Viability of cocultures K0+A0 and KEvo+AEvo inoculated at varying cell densities. The increment in optical density after 72 h of coculturing is depicted in the vertical axis. In panels a, b and c, the average of three replicates is shown. The shaded area and error bars represent standard deviation. In panels d and e, the error bars depict standard deviation across triplicates.

We also observed that, despite a 2.7-fold improvement in the co-culture-level growth rate (**Fig. 3 c**), no significant changes were observable at a single-strain level when they were grown in monocultures (**Fig. 3 a, b**). This suggests that the adaptations resulting from experimental evolution did not improve the overall fitness of either strain, but rather in the specific nutritional and ecological context of a cross-feeding co-culture. To test this, we grew monocultures in low or limiting concentrations of lysine or adenine (**Fig. 3 d, e**), and we observed that the evolved strains KEvo and AEvo grew at higher rates than their ancestral counterparts.

We hypothesised that this improved performance under limited access to cross-fed metabolites could also expand the viable parameter space of the co-culture, allowing it to proliferate from lower inoculation cell densities. This is a known constraint for the viability of co-cultures ^19,20^, which is a strong limiting factor in their applicability in industrial setups. Indeed, our results show a 50-fold reduction in the minimum inoculation cell density, proliferating from concentrations as low as 10 cells/μL (**Fig. 3 f**).

### Modular bioproduction is stabilised by cross-feeding and enhanced by adaptive evolution

To study how the changes triggered by ALE in the ecological properties of the consortium can impact biotechnological applications, we implemented a proof-of-concept split bioproduction pathway. We designed two separate modules with a cross-fed intermediate (**Fig. 4 a**): The first module (PCA module) was engineered to overproduce and secrete p-coumaric acid, a precursor of numerous added-value compounds, while the second module was designed to convert p-coumaric acid into the high-value antioxidant resveratrol (RES module). The PCA module expressed feedback-resistant alleles of three endogenous genes to increase the production of aromatic amino acids (*ARO3*^K222L^, *ARO4*^K229L^ and *ARO7*^G141S^), as well as heterologous genes for p-coumaric acid production via tyrosine and phenylalanine (*PAL* from *Vitis vinifera, C4H* from *Arabidopsis thaliana* and *TAL* from *Flavobacterium johnsoniae*). The second module (RES module) contained genes for the conversion of p-coumaric acid into resveratrol: *4Cl* from *A. thaliana* and *VST1* from *Vitis vinifera*. Coding sequences were expressed from promoter combinations tested in ^31^.

**Figure 4.**
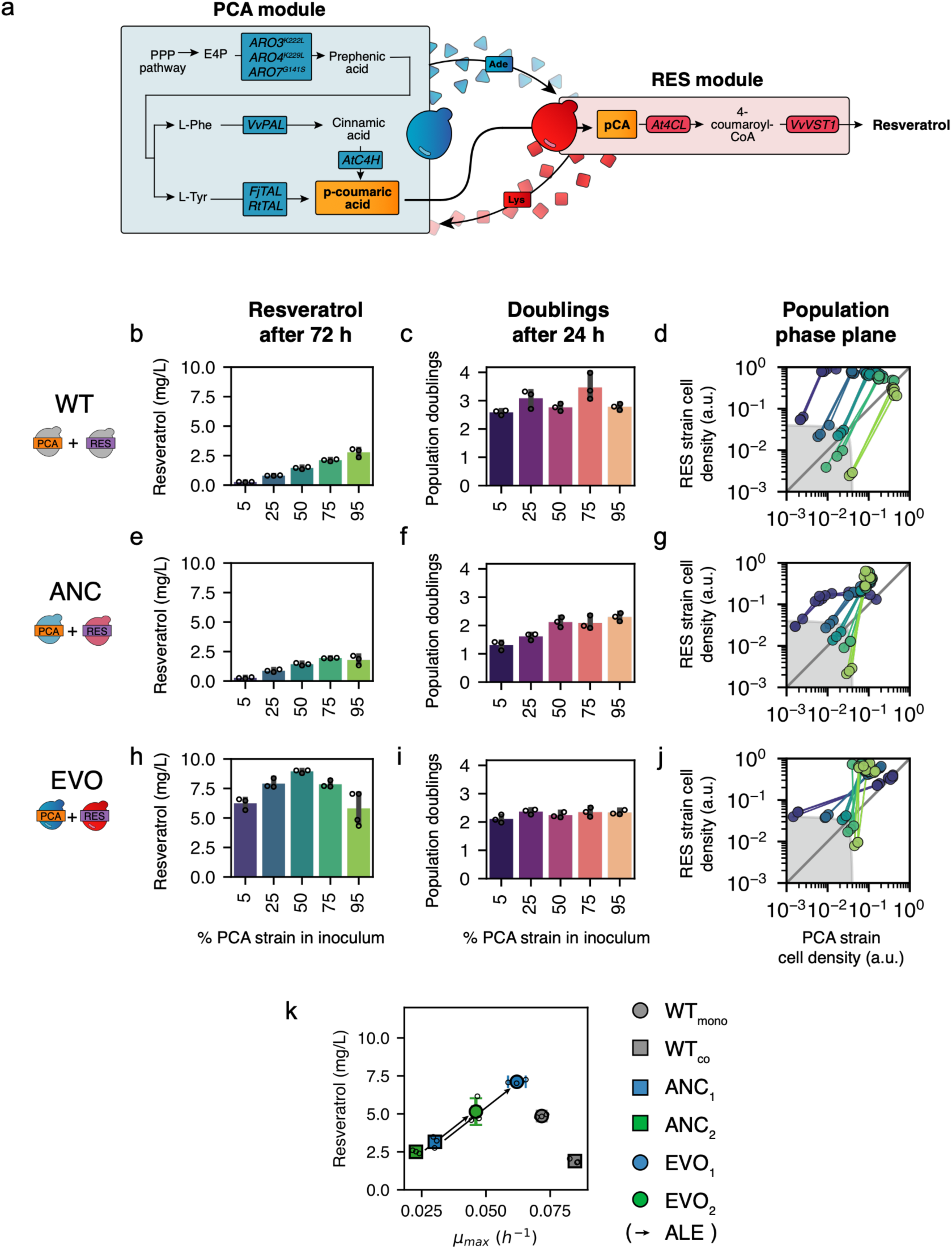
Modular bioproduction in non-cross-feeding, cross-feeding and evolved co-cultures. **a** Resveratrol production pathway split into modules PCA and RES. Coloured boxes depict overexpressed genes: *ARO3*^K222L^, *ARO4*^K229L^ and *ARO7*^G141S^ are feedback-resistant alleles of *Saccharomyces’* endogenous genes, while all other genes are heterologous. Each module is expressed in either strain of the coculture, and the intermediate p-coumaric acid is exchanged across. Adenine and lysine are reciprocally overproduced and exchanged, as described before. **b, e, h** Final resveratrol titre as a function of the starting coculture composition. Strains carrying either the PCA or the RES module were co-inoculated at different relative densities: the horizontal axis depicts the fraction of the inoculum corresponding to the PCA-bearing strain, with the RES-bearing strain making up the reminder. PCA and RES modules were transformed in three co-culture chassis: **b**, “WT”, a co-culture between prototrophic strains, WT::PCA and WT::RES; **e**, “ANC”, a co-culture between K0::PCA and A0::RES; and **h**, “EVO”, a co-culture between KEvo::PCA and AEvo::RES. Bars depict end-point resveratrol titres 72 h after inoculation. Error bars show standard deviation across 3 replicates. **c, f, i** Number of population doublings in the first 24 h after inoculation in the co-cultures and inoculation regimes described in panels b, e and h. **c** is “WT”, **f** is “ANC” and **i** is “EVO”. Error bars show standard deviation across 3 replicates. **d, g, j** Population composition trajectories of the same “WT” (**d**), “ANC” (**g**) and “EVO” (**j**) co-cultures described before, with the same co-inoculation regimes. Each data point represents the absolute abundance of either strain in a given co-culture at a given timepoint. Straight lines connect data points from the same co-culture, sampled every 24 h. The inoculation composition of each co-culture can be inferred from its position in the inoculation space (shaded grey) and the colour scale, which is coherent with panels b and c. Triplicates of each co-culture at each inoculation regime are shown as independent datapoints. **k** Maximum growth rate (“x” axis) and total resveratrol titre after 72 h (“y” axis) in bioproduction co-cultures inoculated at a 1:1 ratio. Arrows connect equivalent co-culture configurations before and after ALE. WTmono is a prototrophic monoculture bearing both PCA and RES production modules; WTco is a co-culture between two prototrophic strains, each bearing one production module; ANC1 is the co-culture K0::PCA+A0::RES; ANC2 is A0::PCA+K0::RES; EVO1 is KEvo::PCA+AEvo::PCA; EVO2 is AEvo::PCA+KEvo::RES. Each marker shows the average of three replicates, depicted as overlaid individual data points. Error bars show standard deviation in each dimension.

The PCA and the RES modules were implemented in three two-strain co-cultures: a non-cross-feeding co-culture composed of two prototrophic strains (WT); the ancestral *ade:lys* co-culture (ANC), composed of K0 transformed with the PCA module and A0 transformed with the RES module; and the evolved co-culture (EVO), composed of KEvo transformed with PCA and AEvo transformed with RES. We co-inoculated these in a range of initial strain ratios, and we measured the final resveratrol titre, growth in the first 24 h, and changes in absolute population composition.

In the WT background (**Fig. 4 b–d)**, resveratrol production was highly sensitive to changes in the initial abundance of the PCA-bearing strain. It ranged from 0.3 mg/L with 5% of the PCA strain in the inoculum, to 2.8 mg/L with 95% (**Fig. 4 b**). The population composition trajectories of this WT co-culture (**Fig. 4 d**) revealed that the PCA strain was strongly outcompeted by the RES strain. This had a particularly noticeable effect at the lower PCA inoculation abundances, where the RES strain was up to 80 times more abundant than the PCA strain after 72 h of growth.

In the ancestral form (ANC, **Fig. 4 e–g**), introducing adenine-lysine cross-feeding considerably dampened the competitive exclusion of the PCA strain: the RES strain was only 1.7 times more abundant than the PCA strain after 72 h, and all population trajectories converged to a central region, regardless of their inoculation composition (**Fig. 4 g**). This convergent behaviour, however, did not result in a weaker dependence of resveratrol production on the inoculation ratio: final resveratrol titres were still as low as 0.3 mg/L at the lowest PCA strain inoculation ratio (**Fig. 4 e**). Interestingly, co-culture level growth also suffered a significant slowdown at those inoculation ratios in early growth phases (**Fig. 4 f**), which was not observed in the WT co-culture (**Fig. 4 c**).

Finally, we studied production in the evolved co-culture (EVO) (**Fig 4. h–j**). Resveratrol titres increased between 23 and 3 times compared to ANC, depending on the co-inoculation ratio, and reached maximum titres at a 50:50 regime (**Fig. 4 h**). In these co-cultures, the convergence of population trajectories observed in ANC was retained (**Fig. 4 j**), but not the growth deceleration at low PCA inoculation ratios (**Fig. 4 i**).

We were interested in comparing our system against a traditional monoculture production setup. We did this in a two-dimensional fitness space showing both production and growth measurements (**Fig. 4 k**). Two configurations of each cross-feeding co-culture were included, where each one of the production modules was transformed in either the adenine- or the lysine-auxotrophic strain (ANC_1_ and ANC_2_; EVO_1_ and EVO_2_). As a monoculture control, we created strain WT_mono_, a prototrophic strain bearing both modules PCA and RES.

First, we compared the behaviour of this prototrophic monoculture implementation (WT_mono_) against a distributed architecture where the production pathway was split across two prototrophic strains (WT_co_). The split pathway design displayed a 17% improvement in growth rate compared to WT_mono_, illustrating how division of labour can redistribute burden associated with the expression of this heterologous pathway. However, resveratrol production was 2.5 times lower in WT_co_: gains along the growth axis came at the expense of performance along the production axis. We next introduced the pathway into the ancestral cross-feeding co-cultures ANC_1_ and ANC_2_. Neither configuration improved production, and both imposed a 2.8- to 3.7-fold growth penalty compared to WT_co_. When we compared these against their equivalent implementation in the evolved chassis (EVO_co1_ and EVO_co2_), we observed that both growth and production increased by 2.2-fold in EVO_1_ over ANC_1_ and 2-fold in EVO_2_ over ANC_2_, even exceeding the production of WT_mono_ in the EVO_1_ configuration by 1.5-fold. These results show that ALE can strengthen the ecological performance of cross-feeding co-cultures and, in doing so, improve production in a heterologous pathway, in some cases exceeding the performance of prototrophic monoculture.

## Discussion

Obligately mutualistic cross-feeding has been extensively implemented in artificial microbial consortia with biotechnological functions. Although its population-stabilising effect is well defined, what role this actually plays in a bioproduction process often remains ambiguous. Furthermore, obligately mutualistic cross-feeding of amino acids and nucleotide bases often results in considerable growth reductions when benchmarked against monocultures in optimal conditions, which can be limiting for their applicability in bioprocesses. Little has been described about the wider implications that such nutritionally restricted contexts can result in. We were interested in studying how adaptive laboratory evolution could be used to improve obligately mutualistic cross-feeding, and whether this could in turn improve production in a co-culture.

We demonstrated that obligate cross-feeding of adenine and lysine can stabilise the population composition of a two-strain *S. cerevisiae* consortia hosting a split bioproduction pathway (**Fig. 4b–g**). The population-stabilising effect of this type of cross-feeding in yeast consortia had been described by Shou *et al*.^19^, but it has only recently been applied in yeast-based bioproduction bioprocesses^20,22^. This is the first time that the population composition trajectories of such a production consortium are explicitly studied, which allowed us to show that, in the absence of obligate cross-feeding, fitness differences result in a progressive exclusion of the precursor-producing strain. This is caused by different levels of metabolic burden imposed by either production module, which is a known issue in the implementation of bioproduction functions in synthetic consortia^16^. We showed that introducing adenine-lysine cross-feeding dampens this effect and guarantees a sustained coexistence of both strains from a range of inoculation ratios. Notably, this original, suboptimal cross-feeding regime did not result in an improvement in resveratrol production, and it considerably reduced co-culture level growth. This highlights how population composition is not the only factor affected by cross-feeding control, as other ecological and physiological processes are altered, and they should be considered in the discussion of artificial cross-feeding for bioproduction.

In fact, transcriptome data (**Fig. 2**) showed how the introduction of lysine and adenine obligately mutualistic cross-feeding profoundly modifies other physiological processes. This consequence of cross-feeding had been described in naturally co-existing microbes^34,35^, but had never been studied in the context of synthetic cross-feeding in a bioproduction platform. We found that both strains experienced a significant slowdown of metabolism, which we hypothesise is an effect of the nutritional restriction imposed by a limited supply of the cross-fed metabolites, as it had been reported before in *Escherichia coli*^36^. Notably, we found a significant activation of Ty1 transposable elements in the adenine auxotrophic strain, which had been described to occur as a result of adenine starvation^33,37^, but never in an obligately-mutualistic co-culture. This is part of a wider stress response linked to purine starvation, which is known to be associated with increased mutation rates^38^ and increased chromosomal instability^39,40^. These observations question the orthogonality of metabolic population control mechanisms and underscore their long-ranging implications in other physiological processes. In fact, the effects we observed are undesired in a bioproduction context, as a reduced general metabolic activity limits productivity, and genetic instability may compromise long bioprocesses.

We described how ALE can be used to address this and improve the performance of obligately mutualistic cross-feeding (**Fig. 1**). Namely, we followed three model co-cultures along a serial passaging experiment, and we found that the progression of this ALE course varied significantly depending on the co-culture. Obligate mutualism is known to impose restrictions on evolutionary trajectories^41^, and in some cases this resulted in abrupt losses or recoveries of viability during the course. Some lineages were taken over by one of the strains, overcoming the obligate dependency, which had been observed in co-cultures that are subjected to stressful conditions^41,42^, further evidencing the physiological strain imposed by suboptimal obligate cross-feeding. Further investigation of these lineages that escaped mutualism could provide valuable insights within the field of evolutionary microbial ecology, but they are out of the scope of this work.

Specifically in the adenine-lysine cross-feeding pair, we found that ALE resulted in significant growth improvements that can be linked with increased uptake and release of the exchanged metabolites (**Fig. 2**). This validated the potential of ALE to engineer such traits in an organism like *S. cerevisiae* ^26,27,29,43,44^, where there is still a big mechanistic knowledge gap in the export of nitrogen metabolites^17,18^. Remarkably, these adaptations lifted the stress response observed in the adenine auxotroph and improved overall metabolic activity across the co-culture, indicating a relief of adenine starvation.

Finally, we demonstrated how general improvements in the physiological state of the co-culture produced by ALE resulted in a significant improvement in resveratrol production (**Fig. 4 h–k**). Interestingly, this was the result of a decoupled evolutionary strategy: ALE was used to improve the overall growth of the co-culture in the absence of a bioproduction function, which was only subsequently introduced. In a production context, we observed how ALE improved the growth restrictions we had previously observed, and improved titres to levels that surpassed production in a non-cross feeding co-culture, but, importantly, also in a prototrophic monoculture. These findings provide a framework for investigating other bioproduction pathways and metabolic families. Future research in this line is also a promising approach to address the stability limitations associated with scaling-up.

Collectively, our results show that the physiological costs associated with obligately mutualistic cross-feeding are not intrinsic limitations, but evolvable traits that can be overcome through ALE. We established a framework for understanding and engineering stable and productive yeast consortia for bioproduction.

## Methods

### Strains and media

All genetic constructs were stored in *Escherichia coli* DH5α and Top10 strains, grown in Luria–Bretani (LB) medium (1% tryptone, 0.5% yeast extract, 1% sodium chloride) at 37 °C or in LB agar with the appropriate antibiotics.

Yeast strains were cloned in a BY4741 background (MATa his3Δ1 leu2Δ0 met15Δ0 ura3Δ0) and grown at 30 °C. Yeast peptone dextrose (YPD) medium was used for transformation, containing 1% yeast extract (Sigma), 2% peptone (Merck) and 2% glucose (VWR). Selection and propagation of transformants was done in SC dropout media formulations, made up of 6.7 g/L yeast nitrogen base without amino acids, 2% glucose and the required yeast synthetic complete (Kaiser) drop-out mixture at the concentration recommended by the manufacturer (Formedium, UK). Minimal medium was prepared with 6.7 g/L yeast nitrogen base without amino acids and 2% glucose. Bacteriological agar (VWR) was added to the media formulations above when preparing plates at 2%. When needed, adenine was supplemented at a final concentration of 10 mg/L in the form of adenine hemisulfate (Sigma) and lysine HCl at 50 mg/L. Unless otherwise stated, co-cultures were inoculated at a total starting OD 600 of 0.8, and monocultures at OD 0.05 for growth and production assays. Yeast strains were stocked in a final concentration of 15% glycerol and stored at -80 °C.

### Plasmid and strain construction

All plasmids used in this study are listed in **Supplementary table 2**. They were built using the MoClo Yeast Toolkit (YTK)^45^ and some backbone sequences from the Multiplex Yeast Toolkit (MYT)^46^. Gene sequences are provided in **Supplementary table 3**, while promoters, terminators and other backbone sequence elements can be found in the YTK publication^45^. Golden Gate was used to assemble these plasmids: 75 fmol of each insert-containing plasmid and 25 fmol of the backbone plasmid were added to a reaction mix containing 0.5 μL T7 ligase (NEB), 0.5 μL type IIS restriction enzyme (BsmbIv2 or BsaI-HFv2, NEB), 1 μL T4 ligase buffer (NEB) and water up to 10 μL. This mix was incubated in a thermocycler with the following programme: 35 two-step cycles consisting of 2 minutes at 42 °C and 5 min at 16 °C, followed by a 10-minute digestion at 42◦C and a 10-minute inactivation at 80◦C. If the enzyme BsaI was used for the assembly, digestion temperature was adjusted to 37◦C. 3 μL of this assembly product was transformed into E. coli cells using a standard heat shock protocol. Assembled plasmids were purified via miniprep using a QIAprep spin miniprep kit (QIAGEN). The correct assembly of intermediate plasmids was verified by restriction digestion, while final plasmids were verified by whole-plasmid sequencing (FullCircle, UK).

Before transforming into yeast, 1 μg of final-construct plasmid was digested with NotI. The lithium acetate protocol was used for transformation^47^: fresh colonies were inoculated from a YPD plate into 3 mL YPD medium and grown to saturation overnight. The following day, they were diluted to an OD 600 nm of 0.5 in 3 mL of YPD and grown up to OD 2. Cells were pelleted and washed twice with sterile water, and once with 0.1 M lithium acetate. Then, they were resuspended in the DNA mix, as well as 240 μL 50% (w/v) PEG, 50 μL 2 mg/mL boiled salmon sperm and 36 μL 1M lithium acetate, and heat-shocked at 42 °C for 40 min. Cells were then pelleted, resuspended in 100 μL sterile water and plated on the appropriate auxotrophic selective medium. Yeast transformation was verified by colony PCR using primers annealing to all relevant coding sequences with the Phire Plat Direct PCR master mix (Thermo). Namely, biomass was boiled for 10 min in a 20 μM sodium hydroxide solution, spun down, and 1 μL of supernatant was used as template for the PCR reaction. PCR was performed according to the Phire Plat Direct PCR master mix manufacturer’s directions and verified in a 1% agarose gel. All strains used in this study are listed in **Supplementary table 1**.

### Adaptive laboratory evolution

Fresh colonies of each co-cultured strain were inoculated in 3 mL of supplemented minimal medium and grown overnight to saturation. Cells were pelleted and washed thrice with sterile water, resuspended in 1 mL of water and their OD 600 nm adjusted. Strains were co-inoculated at a 1:1 ratio in 2 mL minimal medium without supplementation to a final total OD 600 nm of 0.8. in a V-bottom 48 deep-well microplate and sealed with an AeraSeal film (Sigma). These were incubated at 30 °C in a 700 rpm orbital shaking incubator. Every 48 h, cells were transferred to microcentrifuge tubes, spun down at 5,000 rpm in a benchtop centrifuge, washed once with sterile water, and inoculated back in fresh minimal medium at a total OD of 0.8 in a clean deep-well plate. A total of 19 passages were performed, accounting for approximately 75 doublings in the fastest-growing lineages.

### Flow cytometry

The relative abundance of strains in a co-culture was determined by flow cytometry, as strains were differentially tagged by the constitutive expression of a fluorescent protein. All flow cytometry measurements were performed using an Attune NxT Flow Cytometer (Thermo). Samples from liquid cultures were diluted 1:200 in PBS. Cell singlets were gated in the FSC-H/FSC-W and SSC-H/SSC-W planes. In co-cultures, strains were tagged with a constitutively expressed fluorescent protein (mTagBFP2, mScarlet-I or sfGFP), which allowed gating of each strain in the VL1-H, BL1-H or YL2-H channels. Data were collected from at least 10,000 cells in each sample.

### Growth benchmarking

Unless otherwise stated, all growth curves, growth rate measurements and final OD values were collected in plate reader time-lapse experiments. These were performed in a Tecan SPARK or a Biotek Synergy plate reader at the maximum shaking speed for each equipment, double-orbital shaking, with measurements every 30 min. Optical density data were blanked and smoothed by rolling average with a window of 3-5 hours. Smoothed data were used to calculate the instantaneous growth rate at each time interval and the maximum value was extracted. For these experiments, cells were inoculated in 96-well microplates (flat bottom, Costar) in 125 μL of medium and sealed with a Breathe-Easy membrane (Sigma). Monocultures were inoculated at a starting OD of 0.05, while co-cultures were inoculated at OD 0.8.

To test the minimal inoculation density of co-cultures, a master mix of each co-culture was prepared at a 1:1 inoculation ratio and diluted to the desired inoculation densities. Cell concentrations were measured using the Attune Flow NxT cytometer and adjusted accordingly. Co-cultures were inoculated in 2 mL minimal medium in a 48-well deep-well plate sealed with an AeraSeal film (Sigma) and incubated at 30 °C in a 700 rpm orbital shaker for 72 h. After 72 h 100 μL were transferred from each well to a 96-well plate microplate and read in a Byonoy Absorbance 96 microplate reader.

### Resveratrol production assays and population tracking

Strains were co-inoculated in 48-well deep-well plates sealed with an AeraSeal film (Sigma) and incubated at 30 °C in a 700 rpm orbital shaker. Resveratrol production titres were measured after 72 h. Inoculation densities were adjusted using the Attune Flow NxT cytometer cell counts to a total starting density of 4,000 cells/μL and the desired strain ratio. To calculate the doublings after 24 h, 100 μL-samples were taken from the deep-well plate and measured in a Byonoy Absorbance 96 plate reader. To define population trajectories in the phase plane, 10 μL-samples were taken from the deep-well plate every 24 h, diluted and measured by flow cytometry. Absolute cell density measurements are directly provided by the Attune Flow NxT cytometer.

### Metabolite quantification

500 μL of cell culture were mixed with an equivalent volume of HPLC-grade methanol (Sigma) and vortexed for 10 seconds. The mixture was centrifuged at maximum speed for 5 minutes, and 700 μL of the supernatant were transferred to a clean microcentrifuge tube. This volume was centrifuged a second time at maximum speed for 5 minutes and 500 μL of the supernatant were transferred to a glass vial for analysis.

High performance liquid chromatography (HPLC) was used for p-coumaric acid and resveratrol quantification in extracts prepared as above using a Vanquish Core HPLC system (Thermo) equipped with a Discovery® HS F5-5 (10 cm × 4.6 mm, 5 μm, PFP) column (Supelco) and a UV detector. The column compartment was set at 30 °C, flow rate at 1.2 μL/min, and the injection volume was 10 μL. Resveratrol was detected at 306 nm and p-coumaric acid at 289 nm. Elution was carried out in gradient with mobile phases A (10 mM ammonium formate, pH 3) and B (acetonitrile) as follows: 0.0 min, 85% A; 1.5 min, 85% A; 3.0 min, 80% A; 24.0 min, 55% A; 25.0 min, 30% A; 27.0 min, 30% A; 27.5 min, 85% A; 29.0 min, 85% A. The gradient was linear between time points. Peaks were analysed with the software Chromeleon 7.3.1 CDS SE.

### Transcriptomics

Pre-cultures of each strain were grown in SC-LHU medium overnight, washed three times and inoculated at a starting OD of 0.8. Each strain was inoculated in one side of a Duet co-culture unit (Cerillo Bio) in 800 μL of SM. The Duet units were mounted on a carrier plate, sealed with a Breathe-Easy membrane (Sigma) and incubated in a Tecan Spark plate reader at 30 °C, at maximum shaking speed, double orbital. Growth was tracked in the plate reader, cells were harvested at mid-exponential phase and transferred to a microcentrifuge tube. Cells were centrifuged at 5000 g, the supernatant removed by pipetting, and the cell pellet immediately frozen in dry ice and stored at -80 °C.

Total RNA was extracted from frozen cell pellets using the RiboPure RNA Purification Kit yeast (Thermo). RNA quality was assessed from the 260/280 and 260/230 nm absorbance ratios, its concentration determined with a Qubit broad-range assay (Thermo) and integrity estimated by gel electrophoresis.

Library preparation, sequencing and initial bioinformatic analysis were performed by Novogene UK. In brief, messenger RNA was purified using poly-T magnetic beads and cDNA synthesised using random hexamer primers for the first strand. After adapter ligation and size selection, the library was subjected to Illumina sequencing. Raw reads were processed using the fastp software and aligned to the *S. cerevisiae* reference genome assembly R64 (GCF_000146045.2) using Hisat2. featureCounts was used for PFKM calculation and DESeq2 for differential expression analysis grouping samples in triplicates. Enrichment analysis was performed using the GOATOOLS python package and the Genome Association File from the Saccharomyces Genome Database.

### Whole-genome sequencing

Genomic DNA was extracted from *S. cerevisiae* for whole-genome sequencing using the protocol by Denis *et al*.^48^ Quality of was preliminarily assessed from the 260/280 and 260/230 nm absorbance ratios, dsDNA concentration determined with a Qubit broad-range assay (Thermo) and integrity estimated by gel electrophoresis.

Library preparation, sequencing and initial bioinformatic analysis were performed by Novogene UK. In brief, the genomic DNA sample was fragmented, ligated with adapters and PCR-amplified before Illumina sequencing. Raw data were filtered and QC-checked using Fastp and reads aligned to the *S. cerevisiae* reference genome assembly R64 (GCF_000146045.2) using the BWA software and duplicate reads removed using SAMtools. SNP and InDel variant calling was performed using GATK, while copy-number variants and structural variants were identified by CNVnator and BreakDancer, respectively. ANNOVAR was used for functional annotation of variants.

