## Supplementary information for "Adaptive laboratory evolution of a yeast co-culture chassis for modular bioproduction"

#### Supplementary figures

**Supplementary figure 1.** Maximum OD of each combination of clones isolated from the lys2Δ-ADE4op + ade8Δ-LYS21op co-culture.

**Supplementary figure 2.** Aggregated growth rate values by A strain over all K clones, including K0.

**Supplementary figure 3.** Aggregated growth rate values by K clones, excluding combinations with H0.

**Supplementary figure 4.** ORA for biological process GO terms enriched in the subset of genes that were downregulated exclusively in AEvo compared to A0.

**Supplementary figure 5.** Copy number variation in chromosome V of AEvo.

**Supplementary figure 6.** Fold change of FPKM (normalised RNA-Seq reads) between A0 and AEvo of four selected genes.

**Supplementary figure 7.** Growth of KEvo in minimal medium supplemented with lysine and a second amino acid listed in the horizontal axis.

#### Supplementary tables

**Supplementary table 1**. Strains used in this study.

**Supplementary table 2.** Plasmids used in this study.

**Supplementary table 3.** Gene sequences used in this study.

#### Supplementary figures

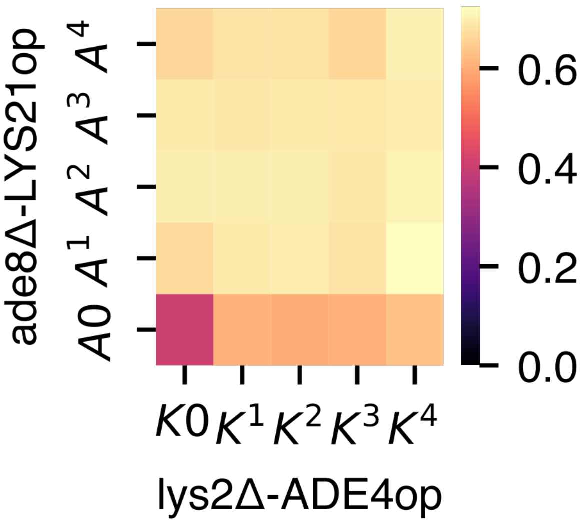


**Supplementary figure 1.** Maximum OD of each combination of clones isolated from the lys2Δ-ADE4op + ade8Δ-LYS21op co-culture. K0 is the ancestral lys2Δ-ADE4op strain, $K^{1\text{-}4}$ are evolved clones of this same strain; A0 is the ancestral ade8Δ-LYS21op strain, $A^{1\text{-}4}$ are its evolved clones. Each tile represents the average value of two replicates.

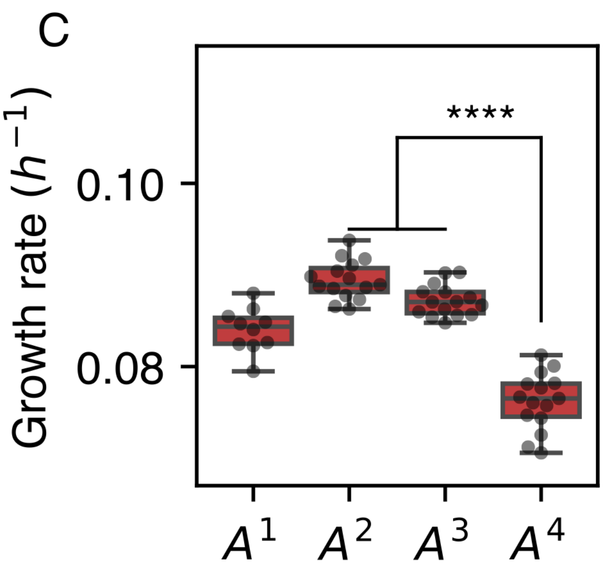


**Supplementary figure 2.** Aggregated growth rate values by A strain over all K clones, including K0. ****: p ≤ 0.0001 in Dunn's test.

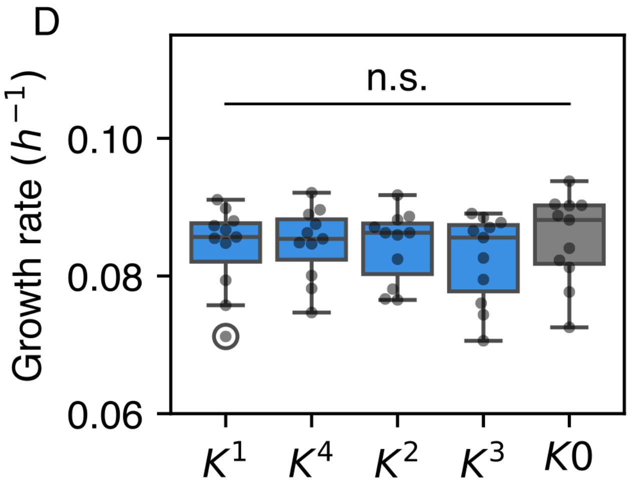


**Supplementary figure 3.** Aggregated growth rate values by K clones, excluding combinations with H0. n.s.: $p > 0.05$ in Kruskal-Wallis test.

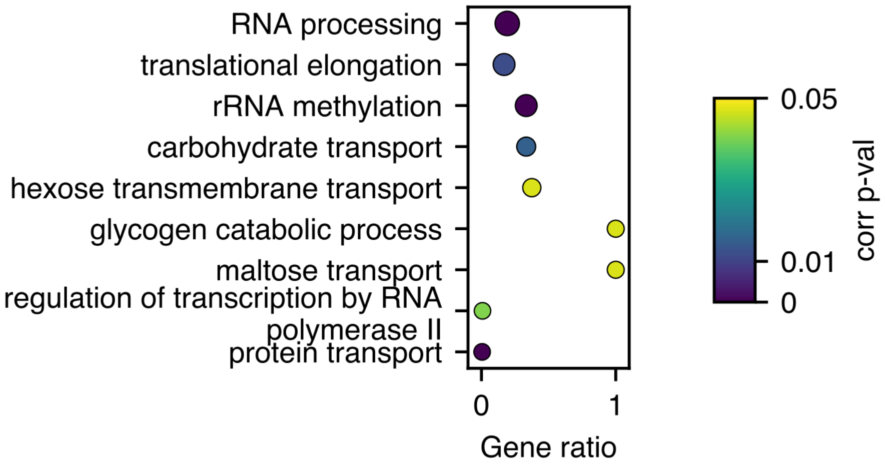


**Supplementary figure 4.** ORA for biological process GO terms enriched in the subset of genes that were downregulated exclusively in AEvo compared to A0. The dot diameter represents gene count, and the colour scale depicts the corrected p-val according to the colour scale in the margin.

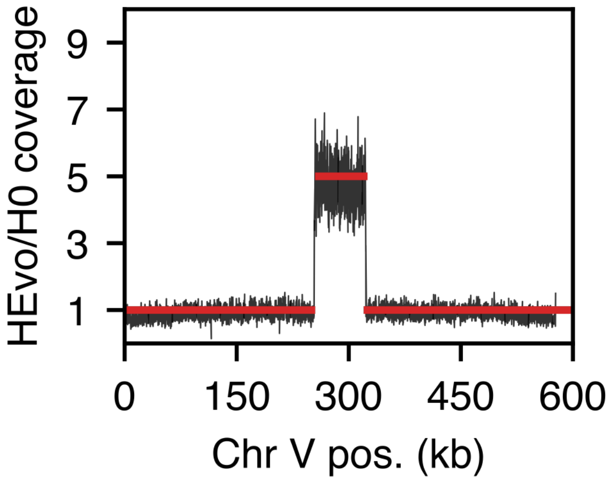


**Supplementary figure 5.** Copy number variation in chromosome V of AEvo. The vertical axis shows the ration between the sequence coverage in A0 and AEvo in short-read WGS, along chromosome V positions (horizontal axis).

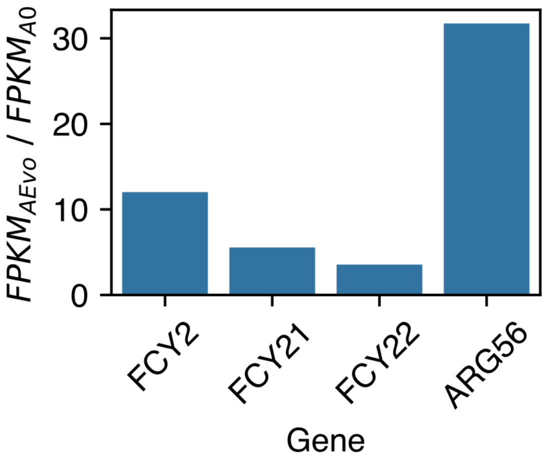


**Supplementary figure 6.** Average fold change of FPKM (normalised RNA-Seq reads) between A0 and AEvo of four selected genes.

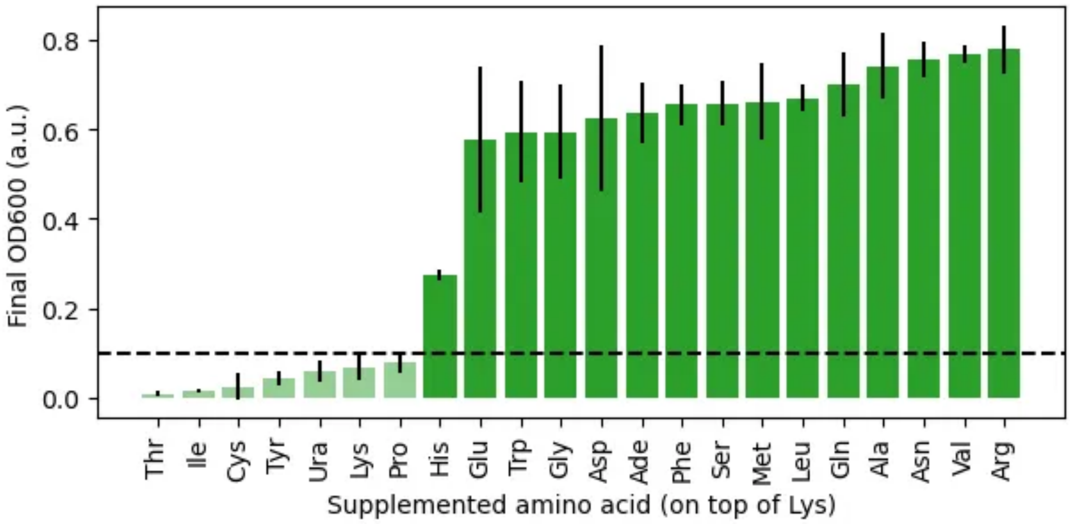


**Supplementary figure 7.** Growth of KEvo in minimal medium supplemented with lysine and a second amino acid listed in the horizontal axis. Error bars show standard deviation across OD 600 after 72 h of growth triplicates.

### Supplementary tables

#### **Supplementary table 1**. Strains used in this study.

| Name | sRLA | Genotype | Source | Plasmids |
|---|---|---|---|---|
| **Adaptive laboratory evolution, transcriptomics** | | | | |
| his2Δ | 339 | BY4741-his2Δ-pTDH3-mTagBFP2-tADH1-vLEU2-pHUM | This work | pHUM, pHP035 |
| met3Δ | 334 | BY4741-met3Δ-pTDH3-mScarlet-I-tADH1-vLeu2-pHUM | This work | pHUM, pHP036 |
| lys2Δ-TRP2op | 1569 | BY4741-lys2Δ-trp2Δ-pTDH3-mTagBFP2-tADH1-vLEU2-pRPL18B-TRP2Fbr-tSSA1-vURA3-pHM | Peng et al., 2024 | pHM, pHP044, pHP035 |
| trp1Δ-LYS21op | 1439 | BY4741-trp1Δ-pTDH3-mScarlet-I-tADH1-vLEU2-pRPL18B-LYS21op-tENO2-vURA3-pHM | Peng et al., 2024 | pHM, pHP054, pHP036 |
| K0 (lys2Δ-ADE4op) | 1301 | BY4741-lys2Δ-pRPL18B-ADE4op-tPGK1-vURA3-pTDH3-mTagBFP2-tADH1-vLEU2-pHM | Peng et al., 2024 | pHM, pHP049, pHP035 |
| A0 (ade8Δ-LYS21op) | 1305 | BY4741-ade8Δ-pTEF1-LYS21op-tENO2-vURA3-pTDH3-mScarlet-I-tADH1-vLEU2-pHM | Peng et al., 2024 | pHM, pHP053, pHP036 |
| KEvo | 3411 | Evolved BY4741-lys2Δ-pRPL18B-ADE4op-tPGK1-vURA3-pTDH3-mTagBFP2-tADH1-vLEU2-pHM | This work | pHM, pHP049, pHP035 |
| AEvo | 3425 | Evolved BY4741-ade8Δ-pTEF1-LYS21op-tENO2-vURA3-pTDH3-mScarlet-I-tADH1-vLEU2-pHM | This work | pHM, pHP053, pHP036 |
| Prot | 3457 | BY4741-URA3:URA3-LEU2:LEU2-pHM | This work | pHM |
| **Production** | | | | |
| WT::pCA | 5617 | MYT2::pTEF1-ARO3[K222L]-tENO1-pCCW12-ARO4[K229L]-tSSA1-pPGK1-ARO7[G141S]-tADH1-HIS3, MYT7::pPGK1-FjTAL-tENO1-pCCW12-PAL-tSSA-pTDH3-AtC4H-tADH1-MET17, URA3::URA3, LEU2::mScarlet-I-vLEU2 | This work | pDR043, pDR044, pHP036 |
| WT::RES | 6022 | MYT2::pTEF1-4CL-tSSA1-pTDH3-VvVST1-tADH1-vHis3Int2, LEU2::mTagBFP2-vLEU2, URA3::URA3, MET15:MET15 | This work | pDR051, pHP035 |
| K0::pCA | 5101 | BY4741, lys2Δ, MYT2::pTEF1-ARO3[K222L]-tENO1-pCCW12-ARO4[K229L]-tSSA1-pPGK1-ARO7[G141S]-tADH1-HIS3, MYT7::pPGK1-FjTAL-tENO1-pCCW12-PAL-tSSA-pTDH3-AtC4H-tADH1-MET17, URA3::ADE4op-vURA3, LEU2::mTagBFP2-vLEU2, | This work | pDR043, pDR044, pHP049, pHP035 |
| A0::pCA | 5106 | BY4741, ade8Δ, MYT2::pTEF1-ARO3[K222L]-tENO1-pCCW12-ARO4[K229L]-tSSA1-pPGK1-ARO7[G141S]-tADH1-HIS3, MYT7::pPGK1-FjTAL-tENO1-pCCW12-PAL-tSSA-pTDH3-AtC4H-tADH1-MET17, URA3::LYS21op-vURA3, LEU2::mScarlet-I-vLEU2 | This work | pDR043, pDR044, pHP053, pHP036 |
| KEvo::pCA | 5103 | BY4741, lys2Δ, evolved, MYT2::pTEF1-ARO3[K222L]-tENO1-pCCW12-ARO4[K229L]-tSSA1-pPGK1-ARO7[G141S]-tADH1-HIS3, MYT7::pPGK1-FjTAL-tENO1-pCCW12-PAL-tSSA-pTDH3-AtC4H-tADH1-MET17, URA3::ADE4op-vURA3, LEU2::mTagBFP2-vLEU2 | This work | pDR043, pDR044, pHP049, pHP035 |
| AEvo::pCA | 5108 | BY4741, ade8Δ, evolved, MYT2::pTEF1-ARO3[K222L]-tENO1-pCCW12-ARO4[K229L]-tSSA1-pPGK1-ARO7[G141S]-tADH1-HIS3, MYT7::pPGK1-FjTAL-tENO1-pCCW12-PAL-tSSA-pTDH3-AtC4H- | This work | pDR043, pDR044, pHP053, pHP036 |

| | | | | |
|---|---|---|---|---|
| | | tADH1-MET17, URA3::LYS21op-vURA3, LEU2::mScarlet-I-vLEU2 | | |
| K0::RES | 5990 | BY4741, lys2Δ, MYT2::pTEF1-4CL-tSSA1-pTDH3-VvVST1-tADH1-vHIs3Int2, URA3::ADE4op-vURA3, LEU2::mTagBFP2-vLEU2, MET15:MET15 | This work | pDR051, pHP035 |
| A0::RES | 5991 | BY4741, ade8Δ, MYT2::pTEF1-4CL-tSSA1-pTDH3-VvVST1-tADH1-vHIs3Int2, URA3::LYS21op-vURA3, LEU2::mScarlet-I-vLEU2, MET15:MET15 | This work | pDR051, pHP036 |
| KEvo::RES | 5721 | BY4741, lys2Δ, evolved, MYT2::pTEF1-4CL-tSSA1-pTDH3-VvVST1-tADH1-vHIs3Int2, URA3::ADE4op-vURA3, LEU2::mTagBFP2-vLEU2, MET15:MET15 | This work | pDR051, pHP035 |
| AEvo::RES | 5725 | BY4741, ade8Δ, evolved, MYT2::pTEF1-4CL-tSSA1-pTDH3-VvVST1-tADH1-vHIs3Int2, URA3::LYS21op-vURA3, LEU2::mScarlet-I-vLEU2, MET15:MET15 | This work | pDR051, pHP036 |

**Supplementary table 2.** Plasmids used in this study.

| Name | pRLA | Description | Source |
|---|---|---|---|
| pHP049 | 1572 | pRPL18B-ADE4op-tPGK1-vURA3 | Peng et al., 2024 |
| pHP044 | 1567 | pRPL18B-TRP2Fbr-tSSA1-vURA3 | Peng et al., 2024 |
| pHP054 | 0578 | pRPL18B-LYS21op-tENO2-vURA3 | Peng et al., 2024 |
| pHP053 | 0577 | pTEF1-LYS21op-tENO2-vURA3 | Peng et al., 2024 |
| pHP035 | 1558 | pTDH3-mTagBFP2-tADH1-vLEU2 | Peng et al., 2024 |
| pHP036 | 1559 | pTDH3-mScarlet-I-tADH1-vLEU2 | Peng et al., 2024 |
| pHM | 15 | HIS3-MET15 | Mulleder et al., 2012 |
| pHUM | 19 | HIS3-URA3-MET15 | Mulleder et al., 2012 |
| pDR043 | 3629 | pTEF1-ARO3[K222L]-tENO1-pCCW12-ARO4[K229L]-tSSA1-pPGK1-ARO7[G141S]-tADH1-HIS3-vMYT2 | This work |
| pDR044 | 3630 | pPGK1-FjTAL-tENO1-pCCW12-VvPAL-tSSA-pTDH3-AtC4H-tADH1-MET17-vMYT7 | This work |
| pDR051 | 3683 | pTEF1-4CL-tSSA1-pTDH3-VvVST1-tADH1-vHIS3Int2 | This work |

**Supplementary table 3.** Gene sequences used in this study.

<table>
<tr><th>Gene name</th><th>Description</th><th>Source</th></tr>
<tr><td>VvPAL</td><td>Phenylalanine ammonia-lyase from Vitis vinifera</td><td>Peng et al., 2023</td></tr>
<tr><td colspan="3">ATGGACGCCACCAACTGCCACGGTTCCAACAAGGTCGAGTCCTTCTGTGTGTCTGACCCCCTGAACTGG<br>GGCATGGCCGCTGAGACTCTCAAGGGTTCTCACCTGGACGAGGTCAAGCGAATGGTGGCCGAGTACCG<br>AAAGCCCGTCGTCCGACTGGGCGGCGAGACTCTGACCATTTCTCAGGTGGCTGCTATTGCTGGACGAGA<br>GGGCGACGTCGGTGTGGAGCTGTCTGAGACTGCTCGAGCCGGCGTCAACGCTTCCTCTGAGTGGGTCA<br>TGGAGTCCATGTCTAAGGGCACCGACTCCTACGGTGTGACCACCGGCTTCGGTGCCACCTCTCACCGAC<br>GAACCAAGCAGGGAGGCGCTCTGCAGAAGGAGCTCATTCGATTCCTGAACGCCGGTATCTTCGGAAACG<br>GACGAGAGTCTTGTCACACCCTGCCTCACTCTGCTACCCGAGCTGCTATGCTCGTCCGAATTAACACCCT<br>GCTCCAGGGATACTCCGGCATCCGATTCGAGATTCTGGAGGCTATCACCAAGCTGCTCAACCACAACATT<br>ACCCCCTGTCTGCCCCTCCGAGGAACCGTGACCGCTTCTGGCGACCTGGTGCCTCTCTCTTACATCGCC<br>GGTCTGCTCACCGGACGACCCAACTCCAAGGCTGTCGGTCCCTCTGGAGAGGTCGTGAACGCTGAGGA<br>AGCCTTCAAGATGGCCGGCATCGAGTCCGGTTTCTTCGAGCTGCAGCCCAAGGAAGGCCTGGCTCTCGT<br>CAACGGAACCGCCGTGGGTTCCGGACTCGCTTCTATGGTGCTGTTCGAGACTAACGTCCTGGCCGTGCT<br>CTCCGAGGTCCTCTCTGCTATTTTCGCCGAGGTCATGCAGGGCAAGCCCGAGTTCACCGACCACCTCAC<br>CCACAAGCTGAAGCACCACCCTGGACAGATTGAGGCCGCTGCCATTATGGAGCACATCCTGGACGGCTC<br>CTCTTACGTCAAGGAAGCCAAGAAGCTCCACGAGATGGACCCCCTGCAGAAGCCCAAGCAGGACCGATA<br>CGCTCTCCGAACCTCTCCCCAGTGGCTGGGACCCCAGATCGAGGTCATCCGAGCTTCCACCAAGTCTAT<br>TGAGCGAGAGATCAACTCCGTCAACGACAACCCCCTGATTGACGTGTCTCGAAACAAGGCCCTCCACGG<br>TGGAAACTTCCAGGGAACCCCCATCGGTGTCTCCATGGACAACACCCGACTCGCCATCGCTGCCATTGG<br>CAAGCTGATGTTCGCTCAGTTCTCCGAGCTCGTGAACGACTTCTACAACAACGGACTCCCCTCTAACCTG<br>GCTGGCTCCCGAAACCCCTCTCTGGACTACGGCTTCAAGGGTGCTGAGATTGCTATGGCCTCCTACTGC<br>TCTGAGCTGCAGTTCCTCGCCAACCCCGTCACCAACCACGTGGAGTCCGCTGAGCAGCACAACCAGGAC<br>GTCAACTCTCTGGGCCTGATTTCCTCTCGAAAGACCGCTGAGGCCGTCGACATCCTGAAGCTCATGTCCA<br>CCACCTACCTGGTGGCTCTCTGTCAGGCCATTGACCTGCGACACCTCGAGGAGAACCTGAAGTCCACCG<br>TCAAGAAAACCGTGTCCCACGTGGCCAAGAAAACCCTGACCATCGGAGCTAACGGCGAGCTCCACCCCT<br>CTCGATTCTGCGAGAAGGACCTGCTCAAGGTCGTGGACCGAGAGCACGTGTTCGCCTACATCGACGACC<br>CCTGTTCCGCTACCTACCCCCTGATGCAGAAGGTGCGACAGGTCCTCGTGGAGCACGCCCTGAACAACG<br>GCGAGTCTGAGAAGAACGGTTCCACCTCTATTTTCCAGAAGATCGGCGCCTTCGAGGAAGAGCTCAAGG<br>CTGTCCTGCCCAAGGAAGTCGAGTCCGCTCGAGATGGCGTCGAGTCCGGAAACCCCTCTATTCCCAACC<br>GAATCAAGGAGTGCCGATCTTACCCCCTGTACAAGTTCGTGCGAGAGGAGCTCGGCACCGGTCTGCTCA<br>CCGGAGAGAAGGTCCGATCCCCCGGAGAGGACTTCGACAAGGTGTTCACCGCCATGTGCGAGGGCAAG<br>ATCATTGACCCCCTGCTCGACTGTCTGTCTGCTTGGAACGGAGCTCCCCTGCCCATCTGTTAATCTAgtgtct<br>gtggtatctaagctatttatcactctttacaacttctacctcaactatctactttaataaatgaatatcgtttattctctatgattactgtatatgcgttcctc</td></tr>
<tr><td>AtC4H</td><td>Phenylalanine ammonia-lyase from Arabidopsis thaliana</td><td>Peng et al., 2023</td></tr>
<tr><td colspan="3">ATGGACCTGCTCCTGCTCGAGAAGTCCCTGATCGCCGTGTTCGTCGCTGTGATTCTGGCCACCGTCATCT<br>CTAAGCTCCGAGGCAAGAAGCTGAAGCTGCCTCCCGGACCCATCCCCATTCCCATCTTCGGAAACTGGC<br>TGCAGGTCGGCGACGACCTGAACCACCGAAACCTCGTGGACTACGCCAAGAAGTTCGGCGACCTCTTCC<br>TGCTCCGAATGGGTCAGCGAAACCTGGTCGTCGTGTCCTCCCCCGACCTCACCAAGGAAGTCCTGCTCA<br>CCCAGGGTGTGGAGTTCGGATCCCGAACCCGAAACGTGGTGTTCGACATTTTCACCGGAAAGGGCCAGG<br>ACATGGTGTTCACCGTGTACGGAGAGCACTGGCGAAAGATGCGACGAATCATGACCGTGCCCTTCTTCA<br>CCAACAAGGTGGTCCAGCAGAACCGAGAGGGCTGGGAGTTCGAGGCCGCTTCCGTGGTCGAGGACGTC<br>AAGAAGAACCCCGACTCTGCCACCAAGGGTATTGTGCTGCGAAAGCGACTGCAGCTCATGATGTACAAC<br>AACATGTTCCGAATCATGTTCGACCGACGATTCGAGTCCGAGGACGACCCCCTGTTCCTCCGACTGAAG<br>GCTCTGAACGGAGAGCGATCCCGACTCGCCCAGTCTTTCGAGTACAACTACGGAGACTTCATTCCCATCC<br>TGCGACCCTTCCTCCGAGGCTACCTGAAGATTTGCCAGGACGTCAAGGACCGACGAATCGCTCTGTTCA<br>AGAAGTACTTCGTGGACGAGCGAAAGCAGATCGCCTCCTCTAAGCCCACCGGATCTGAGGGCCTGAAGT<br>GTGCTATTGACCACATCCTCGAGGCCGAGCAGAAGGGAGAGATTAACGAGGACAACGTCCTGTACATTG<br>TGGAGAACATCAACGTCGCCGCTATCGAGACTACCCTGTGGTCCATTGAGTGGGGCATCGCTGAGCTCG<br>TCAACCACCCCGAGATTCAGTCTAAGCTCCGAAACGAGCTGGACACCGTTCTGGGTCCTGGAGTCCAGG<br>TGACCGAGCCTGACCTCCACAAGCTGCCCTACCTCCAGGCTGTGGTCAAGGAGACTCTCCGACTGCGAA<br>TGGCCATCCCCCTGCTCGTCCCCCACATGAACCTGCACGACGCCAAGCTCGCTGGCTACGACATTCCCG<br>CCGAGTCCAAGATCCTGGTGAACGCTTGGTGGCTCGCCAACAACCCCAACTCTTGGAAGAAGCCCGAGG<br>AGTTCCGACCCGAGCGATTCTTCGAGGAAGAGTCCCACGTCGAGGCTAACGGTAACGACTTCCGATACG<br>TCCCCTTCGGCGTGGGTCGACGATCTTGCCCCGGAATCATTCTCGCCCTGCCCATTCTGGGCATTACCAT<br>CGGTCGAATGGTCCAGAACTTCGAGCTGCTGCCCCCTCCCGGACAGTCCAAGGTGGACACCTCTGAGAA<br>GGGCGGTCAGTTCTCCCTGCACATCCTCAACCACTCTATCATTGTGATGAAGCCCCGAAACTGTTAATCT<br>Agtgtctgtggtatctaagctatttatcactctttacaacttctacctcaactatctactttaataaatgaatatcgtttattctctatgattactgtatatgcgttcctc</td></tr>
</table>

<table>
<tr><td>FjTAL</td><td>Tyrosine ammonia-lyase from Flavobacterium johnsoniae</td><td>Peng et al., 2023</td></tr>
<tr><td colspan="3">ATGAACACCATCAACGAGTACCTGTCTCTGGAAGAGTTCGAGGCCATCATCTTCGGCAACCAGAAGGTGA<br>CCATCTCTGACGTGGTGGTGAACCGAGTGAACGAGTCTTTCAACTTCCTGAAGGAATTCTCTGGCAACAA<br>GGTGATCTACGGCGTGAACACCGGCTTCGGCCCCATGGCTCAGTACCGAATCAAGGAATCTGACCAGAT<br>CCAGCTGCAGTACAACCTGATCCGATCTCACTCTTCTGGCACCGGCAAGCCTCTGTCTCCCGTGTGCGC<br>CAAGGCCGCCATTCTGGCCCGACTGAACACCCTGTCGCTGGGCAACTCTGGCGTGCACCCCTCTGTGAT<br>CAACCTGATGTCTGAGCTGATCAACAAGGACATTACCCCTCTGATCTTCGAGCACGGCGGCGTGGGCGC<br>CTCTGGCGACCTGGTGCAGCTGTCTCACCTGGCTCTGGTGCTGATCGGCGAGGGCGAAGTGTTCTACAA<br>GGGCGAGCGACGACCCACTCCTGAGGTGTTCGAGATCGAGGGACTGAAGCCCATCCAGGTCGAGATCC<br>GAGAGGGACTCGCCCTGATCAACGGCACCTCCGTGATGACCGGCATCGGCGTGGTGAACGTGTACCAC<br>GCCAAGAAGCTGCTGGACTGGTCCCTGAAGTCCTCTTGCGCCATTAACGAGCTGGTGCAGGCCTACGAC<br>GACCACTTCTCTGCCGAGCTGAACCAGACCAAGCGACACAAGGGCCAGCAAGAGATCGCCCTGAAGATG<br>CGACAGAACCTGTCTGACTCTACCCTGATTCGAAAGCGAGAGGACCACCTGTACTCTGGCGAGAACACC<br>GAGGAAATCTTCAAGGAAAAGGTGCAAGAGTACTACTCTCTCCGATGCGTGCCCCAGATTCTGGGCCCC<br>GTGCTGGAAACCATCAACAACGTGGCCTCTATTCTCGAGGACGAGTTCAACTCTGCCAACGACAACCCCA<br>TCATCGACGTGAAGAACCAGCACGTCTACCACGGCGGCAACTTCCACGGCGACTACATCTCCCTCGAGA<br>TGGACAAGCTGAAGATCGTGATCACCAAGCTGACCATGCTGGCCGAGCGACAGCTGAACTACCTGCTGA<br>ACTCTAAGATCAACGAGCTGCTGCCTCCTTTCGTGAACCTGGGCACCCTGGGCTTCAACTTCGGCATGCA<br>GGGCGTGCAGTTCACCGCCACCTCTACCACCGCCGAGTCTCAGATGCTGTCTAACCCCATGTACGTGCA<br>CTCTATCCCCAACAACAACGATAACCAGGACATCGTGTCTATGGGCACCAACTCCGCCGTGATTACCTCT<br>AAGGTGATCGAGAACGCCTTCGAGGTGCTGGCCATCGAGATGATCACCATCGTGCAGGCCATTGACTAC<br>CTGGGCCAGAAGGACAAGATCTCTTCTGTGTCTAAGAAGTGGTACGACGAGATTCGAAACATCATCCCCA<br>CCTTTAAGGAAGATCAGGTGATGTACCCCTTCGTGCAGAAGGTCAAGGACCATCTGATTAACAAC</td></tr>
<tr><td>ARO3-K222L</td><td>DAHP synthase, feedback-resistant</td><td>Peng et al., 2023</td></tr>
<tr><td colspan="3">ATGTTCATTAAAAACGATCACGCCGGTGACAGGAAACGCTTGGAAGACTGGAGAATCAAAGGTTATGATC<br>CATTAACCCCTCCAGATCTGCTTCAACATGAATTTCCAATTTCAGCCAAAGGTGAGGAAAACATTATCAAG<br>GCAAGAGACTCCGTCTGTGATATTTTGAATGGTAAAGATGATCGTTTAGTTATCGTGATCGGGCCATGTTC<br>CCTACATGACCCCAAAGCCGCTTACGATTACGCTGACAGATTGGCTAAAATTTCAGAAAAGTTGTCAAAAG<br>ACTTATTGATTATTATGAGAGCGTATTTAGAAAAACCAAGGACTACTGTTGGCTGGAAAGGGTTGATTAAC<br>GACCCTGATATGAATAACTCTTTTCAAATCAATAAAGGTCTACGGATTTCGAGAGAAATGTTCATAAAACTG<br>GTTGAAAAATTACCCATTGCTGGTGAGATGTTGGATACCATTTCTCCGCAGTTTTTGAGTGATTGTTTCTC<br>CTTGGGTGCCATCGGCGCCAGAACTACTGAATCCCAACTGCACAGAGAATTAGCATCCGGTCTATCTTTC<br>CCTATTGGATTTAAGAACGGTACTGATGGTGGTTTGCAAGTCGCCATCGACGCTATGAGAGCCGCTGCAC<br>ATGAACATTACTTCCTTTCTGTCACATTGCCAGGTGTCACTGCTATCGTGGGCACTGAAGGTAACAAGGAT<br>ACCTTCCTGATCTTGAGAGGTGGTAAGAACGGTACTAACTTTGACAAAGAAAGTGTTCAAAATACTAAGAA<br>ACAGTTAGAAAAGGCCGGTTTGACTGATGATTCCCAGAAAAGAATTATGATCGATTGTTCCCACGGCAAC<br>AGTAATAAAGATTTCAAGAACCAACCAAAGGTTGCCAAATGTATTTATGACCAGCTGACGGAGGGTGAGA<br>ATAGTCTCTGTGGTGTTATGATTGAGTCCAACATAAATGAAGGTAGACAAGATATTCCCAAAGAAGGTGGC<br>AGAGAGGGATTGAAGTATGGTTGTTCTGTTACGGATGCTTGTATTGGCTGGGAGTCCACCGAACAGGTAT<br>TGGAGCTATTGGCAGAAGGTGTTAGAAACAGAAGAAAGGCCTTGAAAAAA</td></tr>
<tr><td>ARO4-K229L</td><td>DAHP synthase, feedback-resistant</td><td>Peng et al., 2023</td></tr>
<tr><td colspan="3">ATGAGTGAATCTCCAATGTTCGCTGCCAACGGCATGCCAAAGGTAAATCAAGGTGCTGAAGAAGATGTCA<br>GAATTTTAGGTTACGACCCATTAGCTTCTCCAGCTCTCCTTCAAGTGCAAATCCCAGCCACACCAACTTCT<br>TTGGAAACTGCCAAGAGAGGTAGAAGAGAAGCTATAGATATTATTACCGGTAAAGACGACAGAGTTCTTG<br>TCATTGTCGGTCCTTGTTCCATCCATGATCTAGAAGCCGCTCAAGAATACGCTTTGAGATTAAAGAAATTG<br>TCAGATGAATTAAAAGGTGATTTATCCATCATTATGAGAGCATACTTGGAGAAGCCAAGAACAACCGTCGG<br>CTGGAAAGGTCTAATTAATGACCCTGATGTTAACAACACTTTCAACATCAACAAGGGTTTGCAATCCGCTA<br>GACAATTGTTTGTCAACTTGACAAATATCGGTTTGCCAATTGGTTCTGAAATGCTTGATACCATTTCTCCTC<br>AATACTTGGCTGATTTGGTTTCCTTCGGTGCCATTGGTGCCAGAACCACCGAATCTCAACTGCACAGAGA<br>ATTGGCCTCCGGTTTGTCTTTCCCAGTTGGTTTCAAGAACGGTACCGATGGTACCTTAAATGTTGCTGTGG<br>ATGCTTGTCAAGCCGCTGCTCATTCTCACCATTTCATGGGTGTTACTTTGCATGGTGTTGCTGCTATCACC<br>ACTACTAAGGGTAACGAACACTGCTTCGTTATTCTAAGAGGTGGTAAAAAGGGTACCAACTACGACGCTA<br>AGTCCGTTGCAGAAGCTAAGGCTCAATTGCCTGCCGGTTCCAACGGTCTAATGATTGACTACTCTCACGG<br>TAACTCCAATAAGGATTTCAGAAACCAACCAAAGGTCAATGACGTTGTTTGTGAGCAAATCGCTAACGGTG<br>AAAACGCCATTACCGGTGTCATGATTGAATCAAACATCAACGAAGGTAACCAAGGCATCCCAGCCGAAGG<br>TAAAGCCGGCTTGAAATATGGTGTTTCCATCACTGATGCTTGTATAGGTTGGGAAACTACTGAAGACGTCT<br>TGAGGAAATTGGCTGCTGCTGTCAGACAAAGAAGAGAAGTTAACAAGAAA</td></tr>
<tr><td>ARO7-G141S</td><td>Chorismate mutase, feedback-resistant</td><td>Peng et al., 2023</td></tr>
</table>

<table>
<tr><td colspan="3">ATGGATTTCACAAAACCAGAAACTGTTTTAAATCTACAAAATATTAGAGATGAATTAGTTAGAATGGAGGAT<br>TCGATCATCTTCAAATTTATTGAGAGGTCGCATTTCGCCACATGTCCTTCAGTTTATGAGGCAAACCATCC<br>AGGTTTAGAAATTCCGAATTTTAAAGGATCTTTCTTGGATTGGGCTCTTTCAAATCTTGAAATTGCGCATTC<br>TCGCATCAGAAGATTCGAATCACCTGATGAAACTCCCTTCTTTCCTGACAAGATTCAGAAATCATTCTTAC<br>CGAGCATTAACTACCCACAAATTTTGGCGCCTTATGCCCCAGAAGTTAATTACAATGATAAAATAAAAAAA<br>GTTTATATTGAAAAGATTATACCATTAATTTCGAAAAGAGATGGTGATGATAAGAATAACTTCTCTTCTGTT<br>GCCACTAGAGATATAGAATGTTTGCAAAGCTTGAGTAGGAGAATCCACTTTGGCAAGTTTGTTGCTGAAG<br>CCAAGTTCCAATCGGATATCCCGCTATACACAAAGCTGATCAAAAGTAAAGATGTCGAGGGGATAATGAA<br>GAATATCACCAATTCTGCCGTTGAAGAAAAGATTCTAGAAAGATTAACTAAGAAGGCTGAAGTCTATGGTG<br>TGGACCCTACCAACGAGTCAGGTGAAAGAAGGATTACTCCAGAATATTTGGTAAAAATTTATAAGGAAATT<br>GTTATACCTATCACTAAGGAAGTTGAGGTGGAATACTTGCTAAGAAGGTTGGAAGAG</td></tr>
<tr><td>At4Cl</td><td>4-coumaroyl-CoA synthase</td><td>Peng et al., 2024</td></tr>
<tr><td colspan="3">ATGGCTCCCCAAGAGCAGGCCGTGTCTCAGGTGATGGAAAAGCAGTCTAACAACAACAACTCTGACGTG<br>ATCTTCCGATCTAAGCTGCCCGACATCTACATCCCCAACCACCTGTCTCTGCACGACTACATCTTCCAGAA<br>CATCTCTGAGTTCGCCACCAAGCCTTGCCTGATCAACGGCCCCACCGGCCACGTGTACACCTACTCCGA<br>CGTCCACGTGATCTCTCGACAGATCGCCGCCAACTTCCACAAGCTGGGCGTGAACCAGAACGACGTGGT<br>GATGCTGCTGCTGCCCAACTGTCCCGAGTTCGTGCTGTCTTTCCTGGCCGCCTCGTTCCGAGGCGCCAC<br>CGCTACCGCTGCTAACCCATTCTTCACCCCTGCCGAGATCGCCAAGCAGGCCAAGGCCTCTAACACCAA<br>GCTGATCATCACCGAGGCTCGATACGTGGACAAGATCAAGCCCCTGCAGAACGATGACGGCGTGGTGAT<br>CGTGTGCATCGACGACAACGAGTCTGTGCCCATTCCTGAGGGCTGCCTGCGATTCACCGAGCTGACCCA<br>GTCTACCACCGAGGCCTCTGAGGTGATCGACTCCGTCGAGATCTCTCCCGACGACGTTGTGGCTCTGCC<br>CTACTCTTCTGGCACCACCGGACTGCCCAAGGGCGTGATGCTGACCCACAAGGGCCTCGTGACCTCTGT<br>GGCCCAGCAGGTCGACGGCGAGAACCCCAACCTGTACTTCCACTCTGACGACGTGATCCTGTGCGTGCT<br>GCCCATGTTCCACATCTACGCCCTGAACTCTATCATGCTGTGCGGCCTGCGAGTGGGAGCCGCCATCCT<br>GATCATGCCCAAGTTCGAGATCAACCTGCTGCTCGAGCTGATCCAGAGATGCAAGGTGACCGTGGCTCC<br>TATGGTGCCTCCTATCGTGCTGGCCATTGCCAAGTCCTCTGAGACTGAGAAGTACGACCTGTCCTCTATC<br>CGAGTGGTGAAGTCTGGCGCTGCTCCCCTCGGCAAGGAACTCGAGGACGCCGTGAACGCTAAGTTCCC<br>CAACGCCAAGCTCGGACAAGGCTACGGCATGACCGAGGCTGGCCCCGTCCTGGCCATGTCTCTGGGCT<br>TCGCCAAGGAACCCTTTCCAGTCAAGTCCGGCGCCTGCGGCACCGTGGTGCGAAACGCCGAGATGAAG<br>ATCGTGGACCCCGACACCGGCGACTCCCTGTCTCGAAACCAGCCTGGCGAGATCTGCATCCGAGGCCA<br>CCAGATCATGAAGGGCTACCTGAACAACCCCGCTGCCACCGCCGAGACTATCGACAAGGACGGCTGGCT<br>GCACACCGGTGACATCGGCCTGATTGACGACGACGATGAGCTGTTCATTGTGGACCGACTGAAGGAACT<br>GATCAAGTACAAGGGCTTCCAGGTGGCTCCCGCCGAGCTTGAGGCCCTGCTGATCGGACACCCCGACAT<br>CACCGACGTGGCCGTGGTCGCCATGAAGGAAGAGGCCGCTGGCGAGGTGCCCGTGGCCTTCGTGGTCA<br>AGTCTAAGGACTCTGAGCTGTCTGAGGACGACGTCAAGCAGTTCGTGTCTAAGCAGGTCGTGTTCTACAA<br>GCGAATCAACAAGGTGTTCTTCACCGAGTCTATCCCCAAGGCTCCCTCTGGCAAGATCCTGCGAAAGGA<br>CCTGCGAGCCAAGCTGGCTAACGGCCTGGGATCCtaa</td></tr>
<tr><td>VvVST1</td><td>Resveratrol synthease from Vitis vinifera</td><td>Peng et al., 2024</td></tr>
<tr><td colspan="3">ATGGCCTCTGTGGAAGAGTTCCGAAACGCCCAGCGAGCCAAGGGACCCGCCACCATCCTGGCCATCGG<br>CACCGCTACTCCCGACCACTGCGTGTACCAGTCTGACTACGCCGACTACTACTTCCGAGTGACCAAGTCT<br>GAGCACATGACCGAGCTGAAGAAGAAGTTCAACCGAATCTGCGACAAGTCTATGATCAAGAAGCGGTAC<br>ATCCACCTGACCGAGGAAATGCTCGAGGAACACCCCAACATCGGCGCCTACATGGCCCCTTCTCTGAAC<br>ATCCGACAAGAGATCATCACCGCCGAGGTGCCCCGACTGGGCCGAGATGCCGCTCTGAAGGCCCTGAA<br>GGAATGGGGACAGCCCAAGTCCAAGATCACCCACCTGGTGTTCTGCACCACCTCTGGCGTCGAGATGCC<br>CGGTGCCGACTACAAGCTGGCCAACCTGCTGGGCCTCGAGACTTCTGTGCGACGAGTGATGCTGTACCA<br>CCAGGGCTGCTACGCTGGCGGCACCGTGCTGCGAACCGCCAAGGACCTGGCCGAGAACAACGCTGGC<br>GCCCGAGTGCTGGTGGTGTGCTCTGAGATCACCGTGGTGACCTTCCGAGGTCCTTCTGAGGACGCCCTG<br>GACTCTCTGGTCGGACAGGCCCTGTTCGGCGACGGATCTTCTGCCGTGATCGTGGGCTCTGACCCCGAC<br>GTGTCTATCGAGCGACCCCTGTTCCAGCTGGTGTCTGCTGCCCAGACCTTCATTCCCAACTCTGCCGGC<br>GCTATCGCCGGAAACCTGCGAGAGGTGGGCCTGACCTTCCACCTGTGGCCTAACGTGCCCACTCTGATC<br>TCTGAGAACATCGAGAAGTGTCTGACCCAGGCTTTCGACCCTCTGGGAATCTCTGACTGGAACTCTCTGT<br>TCTGGATCGCTCACCCCGGTGGACCCGCTATCCTGGACGCCGTCGAGGCCAAGCTGAACCTGGAAAAG<br>AAGAAGCTGGAAGCTACCCGACACGTGCTGTCTGAGTACGGCAACATGTCCTCTGCCTGCGTGCTGTTC<br>ATTCTGGACGAGATGCGAAAGAAGTCTCTGAAGGGCGAGAAGGCCACCACCGGCGAGGGACTCGACTG<br>GGGAGTGCTGTTCGGCTTCGGACCCGGCCTGACCATCGAGACTGTGGTGCTGCACTCTGTGCCCACCGT<br>GACCAAC</td></tr>
</table>
